# Organized cannabinoid receptor distribution in neurons revealed by super-resolution fluorescence imaging

**DOI:** 10.1101/2020.03.22.002642

**Authors:** Hui Li, Jie Yang, Tian Cuiping, Min Diao, Quan Wang, Simeng Zhao, Shanshan Li, Fangzhi Tan, Tian Hua, Chao-Po Lin, Dylan Deska-Gauthier, Garth Thompson, Ying Zhang, Tong Wang, Wenqing Shui, Zhi-Jie Liu, Guisheng Zhong

## Abstract

G-protein-coupled receptors (GPCRs) play important roles in cellular functions. However, their intracellular organization is largely unknown. Through investigation of the cannabinoid receptor 1 (CB_1_), we discovered periodically repeating clusters of CB1 hotspots within the axons of neurons. We observed these CB_1_ hotspots interact with the membrane-associated periodic skeleton (MPS) forming a complex crucial in the regulation of CB_1_ signaling. Furthermore, we found that CB_1_ hotspot periodicity increased upon CB_1_ agonist application, and these activated CB1 displayed less dynamic movement compared to non-activated CB_1_. Our results suggest that CB_1_ forms periodic hotspots organized by the MPS as a mechanism to increase signaling efficacy when being activated.

---

G-protein-coupled receptors (GPCRs) are a large family of membrane proteins that play important roles in cellular functions by initializing a variety of intracellular processes via neurotransmitters and hormone signaling. To fulfill their function, GPCRs have to interact with intracellular protein complexes and are likely anchored by cellular skeletal structures. Therefore, their membrane organization and relationship with skeleton proteins, which are still largely unknown, would be critical for their cellular function. In addition, it would be important to know if diverse GPCRs share similar structural organizations.

Most GPCRs are expressed at low levels in native tissue. Typically, investigators must overexpress them in cultured cells in order to study their structures and functions. However, the cannabinoid receptor 1 (CB_1_) is one of the highest-expressed GPCRs in the central nervous system, making it feasible to study in its natural state ^1^.

CB_1_ is important for many biological functions such as pain, mood and memory ^2, 3, 4, 5^. Recent structural studies have revealed the isolated atomic arrangement of CB_1_ ^6, 7^. However, these studies have primarily used X-ray crystallography or cryo-electron microscopy; techniques which require high concentrations of well-purified proteins in detergent solution. CB_1_ structure has rarely been studied within cells in combination with signaling proteins, integral membrane proteins, or other membrane-associated proteins. Thus, the CB_1_ cellular structure remains incomplete, limiting our ability to fully exploit its function.

Super-resolution microscopy, an imaging method that overcomes the diffraction limit of conventional microscopy, has led to the discovery of the membrane-associated periodic skeleton (MPS) in the axons of neurons ^8^. This highly ordered submembrane skeletal structure can play many roles in neuronal function, including acting as flexible mechanical support, organizing membrane protein distribution, and development of axons and dendrites. The discovery of the MPS, and the many studies that have characterized cellular structures across different cell types^9, 10^, demonstrate the power of super-resolution imaging for uncovering intracellular structures at the nano-scale. Recently, using a type of super-resolution imaging called STORM, Zhou et al. (2019) proposed a model where CB_1_ forms a periodic pattern when activated. They found a ~190 nm periodic pattern of CB_1_ in cultured hippocampal neurons exclusively under the administration of agonists. A previous study, using a similar imaging technology, also revealed distinct CB_1_ structures across different cell types in the brain, but showed no sign of a periodic pattern ^11^. Thus, the intracellular organization of CB_1_ in neurons remains unclear.

Herein, by employing another type of super-resolution imaging, called stimulated emission depletion (STED), we systematically investigated the nano-structure of CB_1_ and other GPCRs in brain tissues and primary cultured neurons. We then generated a new imaging probe that enabled us to monitor the dynamic position of CB1 in live cultured neurons. Our results revealed a periodic structure of CB_1_ clusters along the axons of inhibitory interneurons. These CB1 clusters were organized into “hotspots” ~190 nm apart. Using dual-color STED imaging and cellular biology techniques, we further demonstrated that the CB_1_ hotspots were associated with the MPS. Moreover, the CB_1_ hotspots presented confined dynamics, which were reduced by receptor activation. Thus, our current studies demonstrate that the ~190 nm periodic structure of cytoskeleton appears to the backbone for intracellular signaling to occur.

## Results

### CB_1_ exhibits semi-periodic hotspots in neurons

To study the structure of CB_1_ *in vivo*, we first demonstrated the specificity of our CB_1_ antibody labelling (Supplementary Fig. 1a-b). We found, in agreement with previous findings, that CB_1_ is mainly distributed in the axons of inhibitory interneurons, especially in those of cholecystokinin (CCK)-positive inhibitory interneurons (Supplementary Fig. 1c-d) ^11^. CB_1_ did express much lower in myelinated axons and in the axons of excitatory neurons (Supplementary Fig. 1e-f).

While we confirmed that CB_1_ was distributed in the axon shaft, the nature of its distribution was unknown. Stimulated emission depletion (STED) imaging is known to be well suited for studying the nano-scale structure of cellular components in fixed preparations ^9, 12^. Therefore, we undertook STED imaging of immunolabeled hippocampal tissues to examine CB_1_ organizations along the axon shaft. CB_1_ displayed hotspots both with and without apparent periodicity within axons of the same axonal segment (Fig. 1a-c). We quantified the degree of periodicity using one-dimensional (1D) auto-correlation analysis by projecting the signals to the longitudinal axis of the axon and calculating the average 1D auto-correlation function over many axon segments ^8, 13^. The 1D auto-correlation amplitude, defined as the average amplitude of the peaks at ~190 nm, quantifies the degree of periodicity of the CB_1_ hotspots ^8, 13^.

**Fig. 1.**
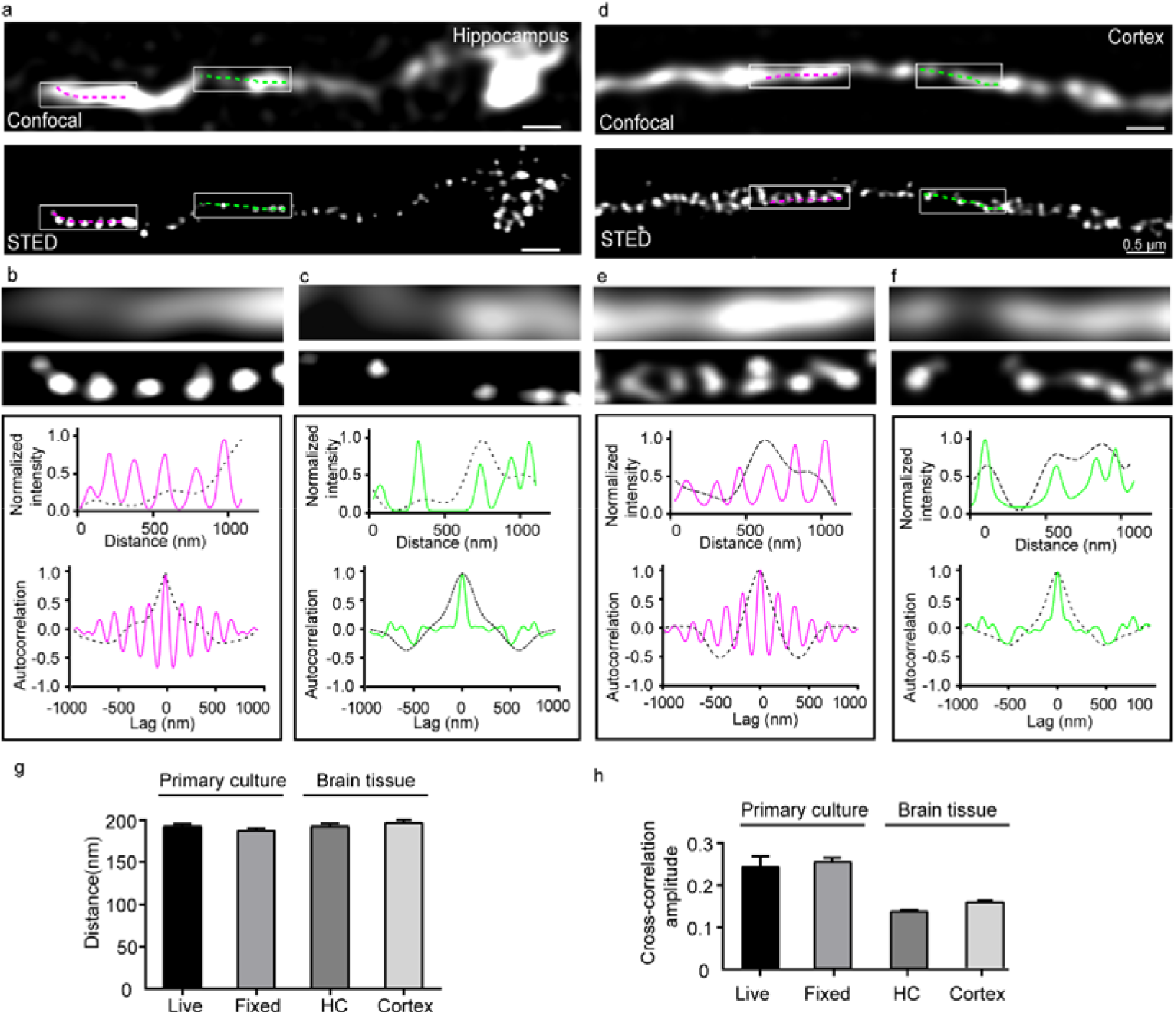
Periodic hotspots of CB_1_ of varying strength in axons from brain tissue. **a** Representative confocal and corresponding STED images of CB_1_ in the hippocampus of mature C57 mice. **b-c** Corresponding magnification of the two boxed regions in **a** (top panel). Intensity plotted along the lines in the box regions (top graph). Autocorrelation analysis for the confocal and STED images (bottom graph). **d-f** Similar to **a-c**, but in the cortex region of mature C57 mice. **g** Histogram of CB_1_ spacing in different samples: live and fixed hippocampal neurons, brain tissue of the hippocampus (HC) and cortex. Data are mean ± s.e.m. (n = 3 biological replicates; 70-120 axonal regions were examined per condition). p = 0.25, (not significant), one way ANOVA. Actual spacing (from left to right): 193 ± 4 nm, 188 ± 5 nm, 189 ± 3 nm, 193 ± 2 nm. **h** Amplitude of the average autocorrelation functions calculated from randomly selected axon segments in different samples. Actual autocorrelation amplitude (from left to right): 0.24 ± 0.03, 0.24 ± 0.05, 0.14 ± 0.07, 0.19 ± 0.05. *p < 0.05 (p = 0.048), one-way ANOVA. Actual p values (from left to right): 0.1534, 0.0125, 0.0239, unpaired Student’s *t* test.

To identify whether CB_1_ possess a similar distribution in other brain regions we performed STED imaging in the cortex, and observed a similar semi-periodicity in neuron axons (Fig. 1d-f). The distance between these rhythmic hotspots was also ~190 nm in the cortex as it was hippocampus (Fig. 1g). Next, we imaged the structure of CB_1_ from cultured hippocampal neurons. We found that antibody-labeled CB_1_ exhibited hotspots with both high and low degree of periodicity in axons of cultured neurons (Supplementary Fig. 2a-c). These results are comparable to the *in vivo* results above (Fig. 1a-c).

To avoid artifacts caused by fixation procedures, we performed live super-resolution imaging on cultured neurons. To this end, we used structured illumination microscopy (SIM), a type of super-resolution imaging that is suitable for investigating the live structure of cellular molecules at high spatial resolution^14^. SIM imaging revealed clusters of CB_1_ in cultured neurons, and those clusters appeared highly organized as hotspots in some regions of axons (Supplementary Fig. 2d-f). Again, CB_1_ exhibited both high and low periodic hotspots (Supplementary Fig. 2d-f). The spatial distance between periodic hotspots was ~190 nm, which was comparable to our results with antibody labelling in fixed preparations (Fig. 1g). Notably, the regularity of the CB_1_ structure in cultured neurons was higher than that in brain (Fig. 1h). Therefore, we conclude that there is a commonality of the semi-periodic feature of CB_1_ in neurons, an unexpected characteristic of GPCRs in the cellular membrane.

### CB_1_ is associated with MPS in neurons

The semi-periodic organization of CB_1_ in axons raised two important questions: how is the CB_1_ semi-periodic organization formed and what are the complexes related to the semi-periodic organization of CB_1_? To answer the first question, we need to identify the components associated with the semi-periodic complex of CB_1_ in native states. To this end, we performed mass spectrometry (MS) experiments with modifications to increase the enrichment of membrane proteins in six different regions of the mouse brain (Supplementary Fig. 3). We found that CB_1_ was expressed at different levels in the six brain regions and that the expression level paralleled the glutamate receptor 1 and N-Methyl-d-aspartate (NMDA) receptor (Supplementary Fig. 3a), which are known to crosstalk with CB_1_ in the central nervous system ^15, 16^. Furthermore, we observed the association of GPCR-related signaling molecules with CB_1_ (such as Gi, tyrosine-protein kinase and Fyn), indicating the reliability of our modified MS method (Supplementary Fig. 3b).

Next, we compared the expression extent of selected MPS components with CB_1_ expression. While different isoforms of ankyrin expressed across different functional domains of neurons, ankyrin-B (ankB) was predominantly distributed in the axons of neurons ^13, 17^. Our MS experiments indicated that the expression level of CB_1_ was highly correlated to that of ankB, but not ankyrin-R (ankR) (Supplementary Fig. 3a). Further, the expression level of CB_1_ was correlated to that of both αII-spectrin and βII-spectrin (Supplementary Fig. 3a) while other membrane proteins seem unrelated to the expression levels of CB_1_ (Supplementary Fig. 3c).

Next, we directly visualized the spatial relation of CB_1_ and MPS molecules by dual colour STED imaging. The MPS was visualized through immunolabelling of the C-terminus of βII-spectrin, which is located at the center of each spectrin tetramer connecting adjacent actin rings ^13, 17^. We quantified the extent of colocalization between CB_1_ and βII-spectrin using 1D cross-correlation analysis to calculate the average 1D cross-correlation function between the two color channels over randomly chosen axon segments ^8^. At high periodic hotspots, CB_1_ displayed high colocalization with βII-spectrin while at non-periodic clusters, CB_1_ displayed little colocalization with βII-spectrin (Fig. 2a-c). Previous studies demonstrated that ankB mediated the attachment of membrane proteins to the MPS, and is located in the middle region of each spectrin tetramer ^17, 21^. Thus, we reasoned that CB_1_’s interaction with ankB may organize the semi-periodic pattern of CB_1_ with the MPS. Once again, at high periodic hotspots, CB_1_ displayed high colocalization with ankB while at non-periodic clusters, CB_1_ displayed little colocalization with ankB (Supplementary Fig. 4a-c). Furthermore, the spatial distance between periodic CB_1_ hotspots was comparable to both those of βII-spectrin and ankB (Supplementary Fig. 4d). These results unambiguously show that CB_1_ forms a semi-periodic complex associated with the MPS.

**Fig. 2.**
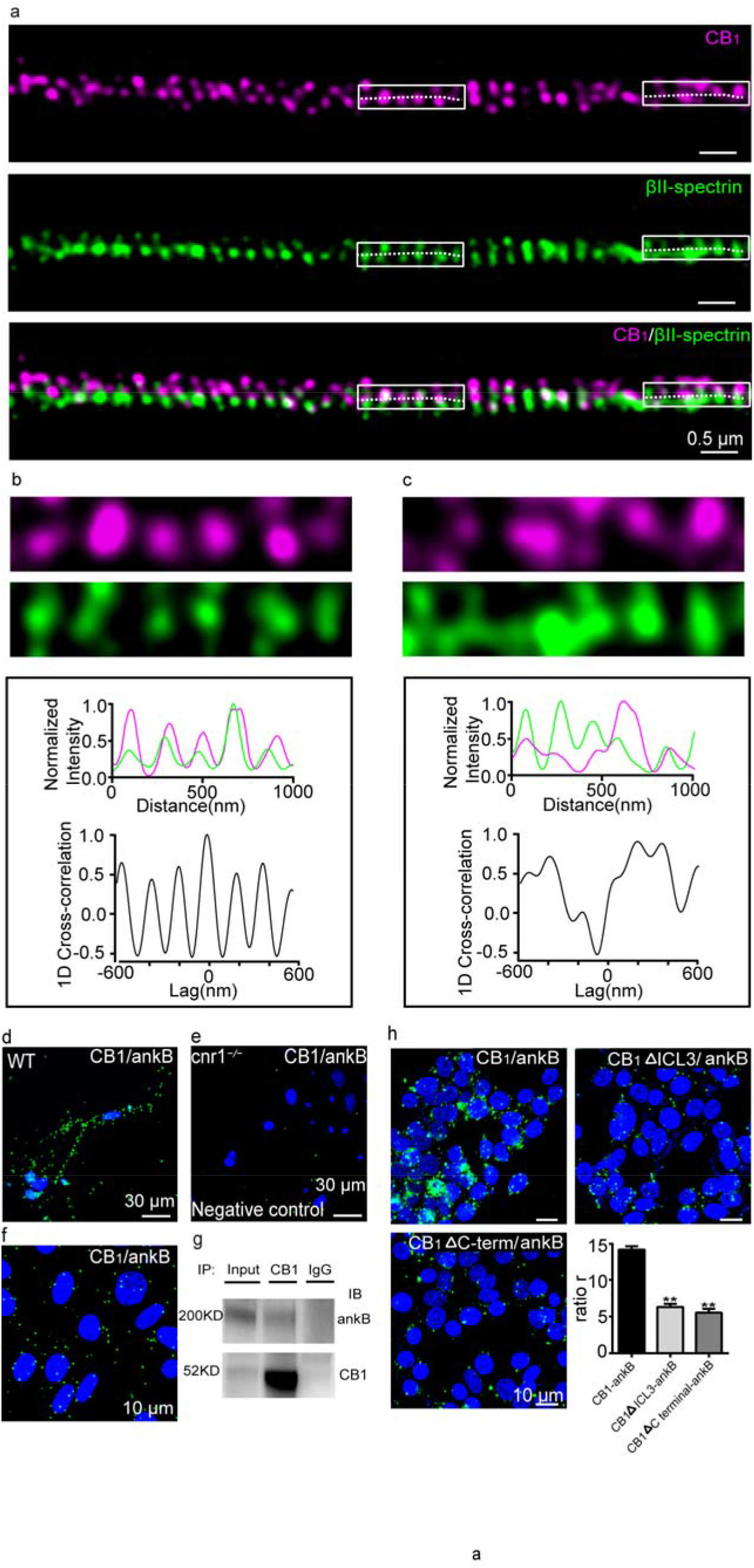
CB_1_ hotspots are connected with components of the MPS. **a.** Two-color STED images of CB_1_ (magenta) and βII-spectrin (green) in the axons of cultured neurons. **b-c.** Top, enlarged images taken from white boxes from **a**. 1D projection traces of βII-spectrin (green) and CB_1_ (magenta) signals along the axon are shown in the middle. 1D cross-correlation functions between the distributions of CB_1_ and βII-spectrin from CB_1_-positive axon segments are shown in the bottom. **d-f.** PLA was performed in cultured neurons of WT mouse **(d)**, *cnr1*^−/−^ mouse **(e)**, and tetracycline-induced CB1 transfected CHO cells (d) with antibody of CB_1_ and ankB. Cell nuclei were stained with DAPI (blue). **g.** Immunoprecipitation of CB_1_ and ankB in CB_1_-CHO cells. Samples were processed for immunoprecipitation with either anti-CB_1_ or IgG control antibodies. Immunoprecipitates were immunoblotted with the anti-CB_1_ antibody (55kD) and anti-ankB antibody (220kD). **h.** PLA was performed in HEK-293T cells co-expressing ankB and different fragments of CB_1_, including wildtype (CB1), CB_1_ with truncated ICL3 loop (CB1ΔICL3), or CB_1_ with truncated C-terminal (CB1ΔC-term). Data are mean±s.e.m. (n=5 biological replicates; 25-50 imaging regions were examined per condition). **p < 0.001, unpaired Student’s *t* test. Actual p values (from left to right): 2.5×10^−4^, 1.6×10^−4^.

To demonstrate the association between CB_1_ and the complex components identified by the MS experiments, we carried out proximity ligation assays (PLA) ^18^, another type of imaging assay that reliably detects the close physical distribution of two subjects. In cultured neurons, we determined that CB_1_ localizes closely with complex components. PLA signals of CB_1_ and ankB were observed in cultured primary neurons (Fig. 2d), but no PLA signals were detected in the *cnr1*^−/−^ mouse neurons (Fig. 2e). It is known that α-adducin and βII-spectrin closely distributed in neurites. Indeed, in *cnr1*^−/−^ mouse, PLA signal with α-adducin and βII-spectrin was not changed (Supplementary Fig. 5a-b), indicating the reliability of PLA to detect the proteins within a short distance. Upon the induction of CB_1_, close physical distances were detected between CB_1_ and several complex components, including ankB, βII-spectrin and αII-spectrin in CB1-CHO cell line (Fig. 2f, Supplementary Fig. 5c-e). These results illustrate the close physical distance between CB_1_ and complex components.

In order to show that CB_1_ interacted with ankB in tetracycline-induced CB_1_-CHO cells, we performed immunoprecipitation experiments (Fig. 2g). To identify which parts of CB_1_ may mediate the interaction between CB_1_ and ankB, we truncated the third intracellular loop (ICL3) or C terminal of CB_1_. Notably, the PLA signals were significantly decreased in both conditions, suggesting that both regions, ICL3 and C-terminal, play a role in the interaction between CB_1_ and ankB (Fig. 2h). Together, these results support the view that CB_1_ may form a complex with receptor kinases and G proteins together with cytoskeleton related proteins, such as ankB, αII-spectrin and βII-spectrin.

### CB_1_ displays confined dynamics in neurons

CB_1_ connects to the MPS through ankB forming semi-periodic hotspots. As such, we would expect CB_1_ to display confined dynamics by interaction with the MPS through ankB. Therefore, we investigated the dynamics of individual hotspots of CB_1_ with live SIM imaging. Neurons were ectopically expressed with a CB_1_-RFP fusion protein (RFP protein is fused to the end of CB_1_ C-terminal) and live SIM images were acquired 1 day after transfection. In order to identify whether CB_1_-RFP was recruited to the native site, we performed two color STED imaging of CB_1_-RFP and βII-spectrin. CB_1_-RFP displayed highly periodic hotspots that co-localized with βII-spectrin, as well as non-periodic clusters that not colocalized with βII-spectrin (Supplementary Fig. 6a-c) These results were similar to the spatial distribution between CB_1_ and βII-spectrin (Fig. 2a-c), supporting that CB_1_-RFP is transported to CB_1_ native sites. We proceeded to examine the dynamics of CB_1_-RFP using live SIM imaging. Periodic CB_1_ hotspots were confined to movements around their starting positions displaying confined displacement changes (average 69 nm) between time frames (Fig. 3a-c). Averaged autocorrelation analysis of CB_1_ at different time frames showed a similar periodic distribution, indicating that CB_1_ clusters maintained their periodicity through time (Fig. 3d). The cross correlation of neighboring time points was calculated that showed the average cross correlation value peaked at zero point (Fig. 3e), indicating little or no systematic shift of the CB_1_ periodicity across different time frames. The periodic wavelength was around 190 nm at different time frames (Fig. 3f). We compiled the moving traces of individual hotspots of CB_1_ that clearly indicated confined movement both at short (2 min) (Fig. 3g) and longer (6 min) (Supplementary Fig. 7a-g) imaging time frames. The moving traces between the two time frames were comparable (Supplementary Fig. 7h), suggesting that CB_1_ displayed a confined movement around its anchoring point, likely mediated by ankB.

**Fig. 3.**
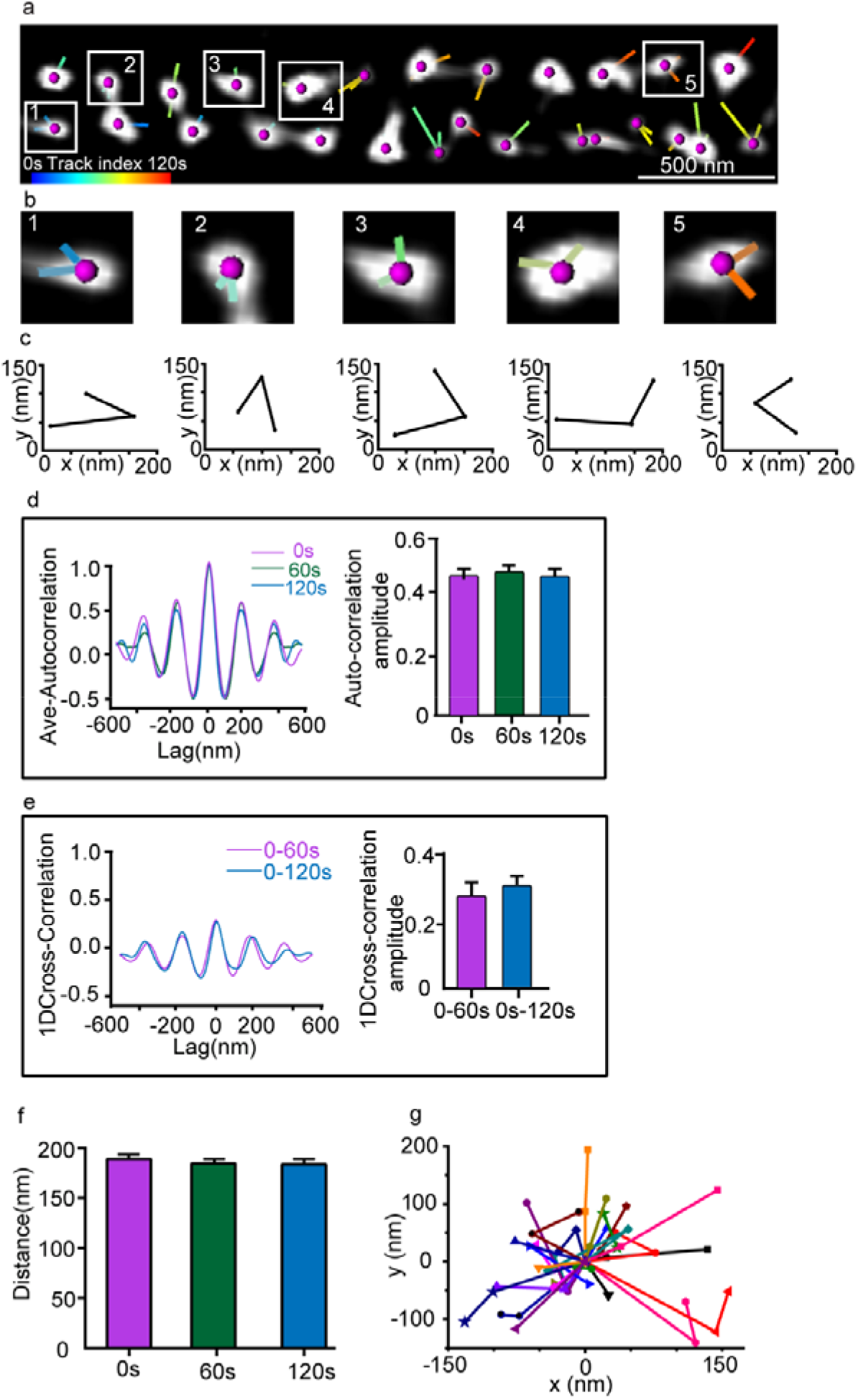
Periodic CB1 hotspots display stable dynamics revealed by live SIM imaging. **a** Representative live image of transfected CB_1_-RFP in the primary neuron (DIV 9-12) of the SD rat acquired by SIM. Individual CB_1_ hotspots are marked with purple balls, and locations of each individual time points are connected with lines. Line colors indicate trace indexes. **b** Five CB_1_ hotspots shown in **a** with their relative locations. **c** Displacement changes from **b** are around 60-70 nm between neighboring time points. **d** Averaged autocorrelation analysis of CB_1_ distributions at different time points with the histogram of the autocorrelation amplitude. p = 0.8757 (not significant), one-way ANOVA. **e** Averaged cross-correlation analysis between the neighboring frames (0s vs 60s, 0s vs 120s) showed similar distribution properties and histogram showing amplitude of average cross-correlation. p = 0.589, (not significant), unpaired Student’s *t* test. Actual cross-correlation amplitude (from left to right): 0.26 ± 0.06; 0.30 ± 0.04. **f** The histogram of CB_1_ spacing across time points. p = 0.4335 (not significant), one-way ANOVA. Actual spacing (from left to right): 192 ± 1 nm, 191 ± 1 nm, 191 ± 1 nm. Data in **d**-**f** are mean ± s.e.m. (n = 3 biological replicates; 70-120 axonal regions were examined per condition). **g** Traces of the individual CB_1_ hotspots over time.

### Active CB_1_ tightly associate with MPS and display less dynamics

We proceeded to examine the dynamics of active CB_1_ upon the CB_1_ agonist, WIN 55,212-2 (WIN), application using live SIM imaging. Previously, several probes have been developed to detect neuromodulator receptor activations, such as dopamine (DA), norepinephrine (NE), and acetylcholine (Ach) ^22, 23, 24^. An agonist-induced conformational change of these GPCRs alters the arrangement of the associated cpGFP and results in a fluorescence change. We took a similar strategy by optimally inserting a conformationally sensitive cpGFP into the third intracellular loop (ICL3) of CB_1_. WIN-induced fluorescence responses were rapidly reversed by the CB_1_ antagonist, rimonabant (Fig. 4a-b). The probe did not respond to ligands of other GPCRs, indicating its specificity (Fig. 4c).

**Fig. 4.**
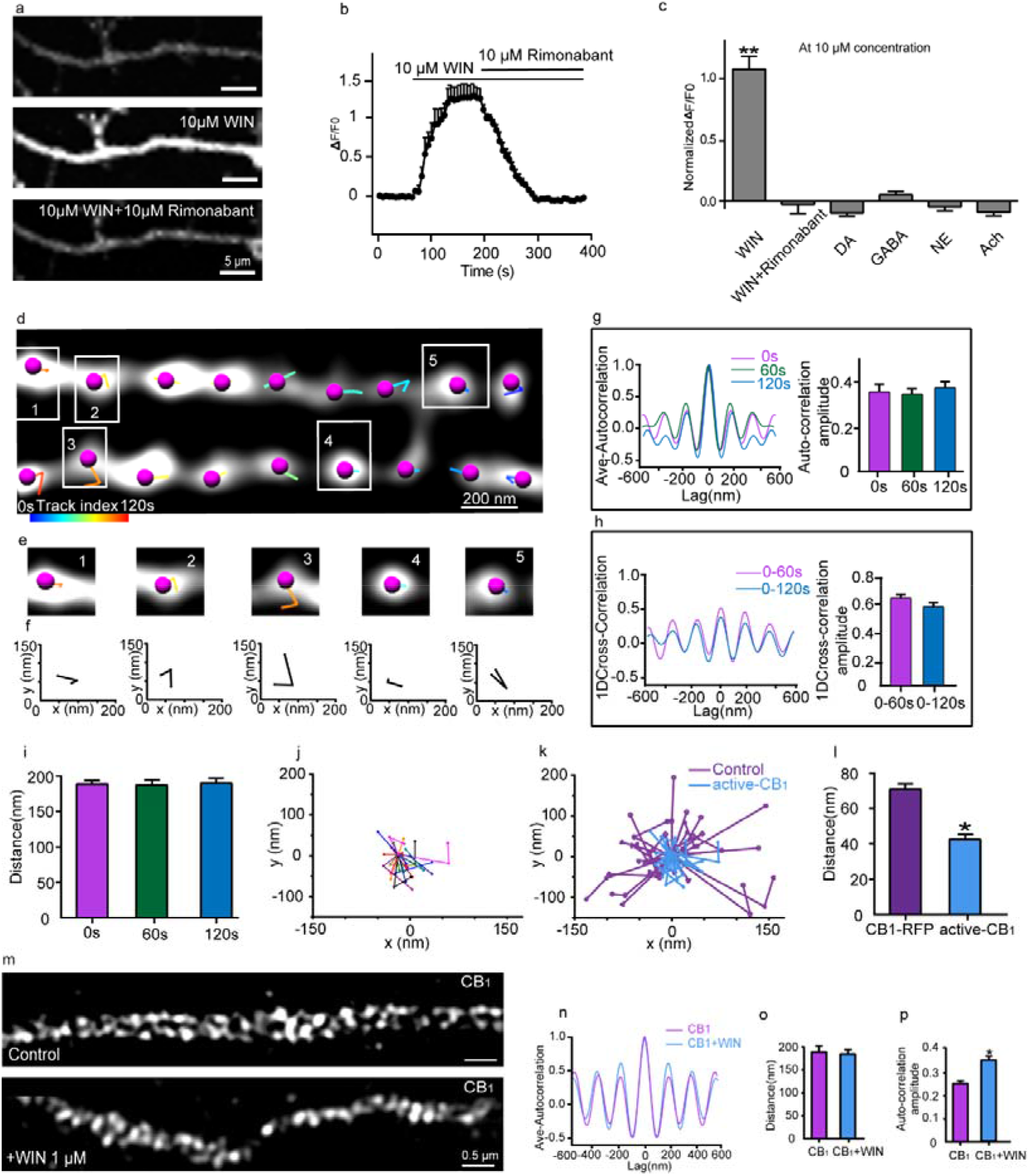
The dynamics of active CB_1_ hotspots revealed by live SIM imaging**. a** Representative confocal images of CB_1_ probe in primary hippocampal neurons with different treatments of CB_1_ ligands. **b** Traces of fluorescence changes in CB_1_ probe-expressing neurons in response to 10 μM WIN followed by 10 μM rimonabant. **c** Normalized fluorescence changes in response to the application of indicated compounds. Actual normalized ΔF/F0 (from left to right): 1 ± 0.099, n = 7; −0.024 ± 0.071, n = 7; −0.076 ± 0.022, n = 9; 0.061 ± 0.025, n = 9; −0.066 ± 0.032, n = 9; −0.082 ± 0.0272, n = 9. *p < 0.0001, n = 3 biological replicates, unpaired Student’s *t* test. Actual p values (from left to right): 7.8×10^−4^, 2.1×10^−4^, 3.6×10^−4^, 4.7×10^−4^, 5.9×10^−4^. **d** Representative live image of active CB_1_ in the primary neuron (DIV 9-12) of the SD rat acquired by SIM. **e** Five active CB_1_ hotspots shown in **d** with their relative locations. **f** Displacement changes from **e** are around 40-50 nm between neighboring time points. **g** Averaged autocorrelation analysis of active CB_1_ distributions at different time points with the histogram of the autocorrelation amplitude. p = 0.1661, not significant (p>0.1), one-way ANOVA. Actual autocorrelation amplitude (from left to right): 0.34 ± 0.02, 0.31 ± 0.03, 0.39 ± 0.03. **h** Averaged cross-correlation analysis between the neighboring frames (0s vs 60s, 0s vs 120s) showed similar distribution properties, and histogram shows amplitude of average cross-correlation. p = 0.7733, (not significant), unpaired Student’s t test. Actual cross-correlation amplitude (from left to right): 0.64 ± 0.05; 0.59 ± 0.16. i Histogram of CB_1_ spacing across time points. p = 0.8504, (not significant), one-way ANOVA. Actual spacing (from left to right): 190 ± 4 nm, 191 ± 5 nm, 192 ± 5 nm. Data in **g-i** are mean ± s.e.m. (n = 50 biological replicates; 70-120 axonal regions were examined per condition). **j** Traces of the individual CB_1_ hotspots over time. **k** Trace the dynamics of CB_1_ (purple) and active CB_1_ (blue) hotspots over time. **l** Actual displacement (from left to right): 69 ± 4, 42 ± 4. n =109, n = 36 spots. p = 0.0002, *p < 0.01, unpaired Student’s t test. **m** Representative SIM images of CB_1_ in primary hippocampal neurons for control and with WIN treatment. **n** Autocorrelation analysis for the SIM images of CB_1_. **o** Histogram of the distance of CB_1_ hotspots for control and with WIN treatment. p = 0.56, (p < 0.5), unpaired Student’s t test. Actual spacing (from left to right): 193 ± 4 nm, 190 ± 3 nm. **p** The histogram of autocorrelation amplitude of CB_1_ without and with WIN treatment was shown. p = 0.04, (*p < 0.05), unpaired Student’s t test. Actual autocorrelation amplitude (from left to right): 0.24 ± 0.03, 0.35 ± 0.04.

Using SIM microscopy, we observed the periodic hotspots of CB_1_ upon WIN application (Fig. 4d). Active CB_1_ hotspots were confined to movements around their starting position between time frames (Fig. 4d-f). Averaged autocorrelation analysis of CB_1_ at different time frames showed similar periodic distribution, indicating the active CB_1_ hotspots maintained their periodicity through time (Fig. 4g). To find out if there was any space shift of such periodicity, the cross correlation of neighboring time points was calculated. The average cross correlation value peaked at the zero point (Fig. 4h), indicating no systematic shift of the active CB_1_ periodicity across different time frames. The periodic space was around 190 nm at different time frames (Fig. 4i). We compiled the moving traces of individual hotspots of CB_1_ (Fig. 4j), clearly indicating a confined movement. Finally, our results showed that with WIN application, active CB_1_ moved around their original point to a lesser extent than non-active CB_1_ (Fig. 4k-l). Next, we treated neurons with the CB_1_ agonist, WIN, and fixed the samples for immunolabelling with CB_1_ antibody. We found that CB_1_ displayed hotspots with higher periodicity when WIN was applied versus the native state (Fig. 4m-p).

### CB_1_ signaling is related to cytoskeleton

To evaluate the dependence of CB_1_ periodic hotspots on the cytoskeleton, we treated cultures with either latrunculinB (latB) or cytochalasin D (cytoD) to disrupt cytoskeletal structure ^8, 13^. We found that following both latB and cytoD treatments, periodic hotspots of CB_1_ were no longer observed (Fig. 5a-b), suggesting that the cytoskeleton is important for maintaining the CB_1_ structure in neurons. Next, we tested whether the cytoskeleton affects CB_1_ intracellular downstream signaling. CB_1_ can activate both the ERK1/2 ^25, 26^. In primary neuron cells, pre-treatment with latB resulted in decreased phosphorylation levels of both Akt and ERK1/2 in a dose-dependent manner (Supplementary Fig. 8a-b). Thus, these data suggest that the intracellular signaling of CB_1_ is dependent upon the MPS cytoskeleton (Fig. 5c-e).

**Fig. 5.**
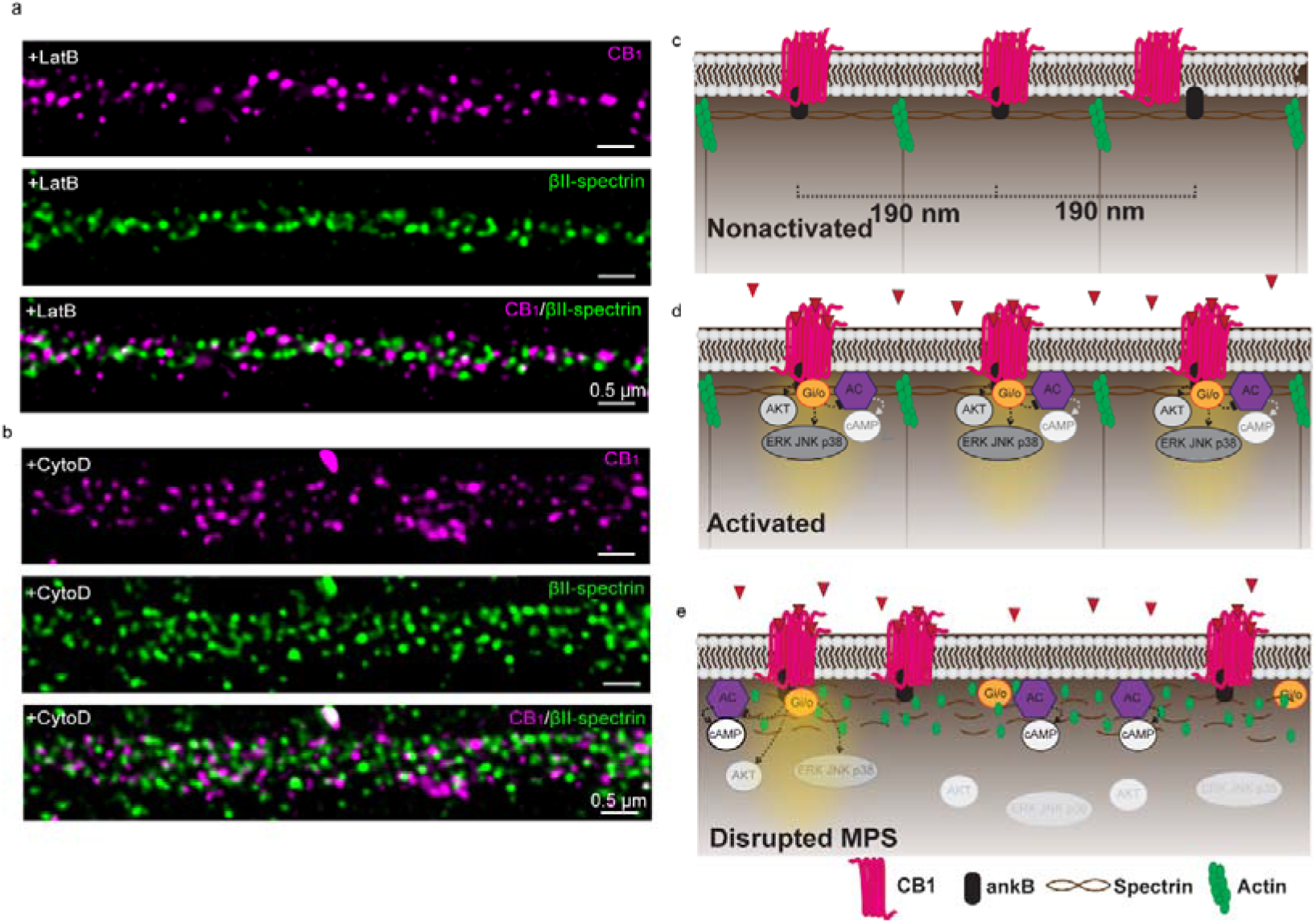
Schematic illustration showing CB_1_ forming dynamic peri-periodic hotspots to increase signaling efficiency. **a-b** Two-color STED images of CB_1_ (magenta) and βII-spectrin (green) in neurons treated with LatB (**a**) and CytoD (**b**). **c** Without ligand binding, MPS sets the range for CB_1_ distribution. **d** Upon ligand binding, active CB_1_ are recruited to the MPS and become more periodic, making downstream AKT and ERK signaling more effective. **e** MPS degradation leads to less strong periodic clusters of CB_1_ and thus less downstream signaling.

## Discussion

Visualization of GPCRs in native tissue is necessary for understanding the intracellular organization of these receptors in real physiology. With the recent development of super-resolution imaging methods, we now can observe the organization of GPCRs at the cellular and tissue levels ^11, 27^. It is also important to visualize the live dynamics of GPCRs in order to understand potential functional changes in physiological and pathological conditions. Here, we used SIM to investigate the dynamics of CB_1_. Our findings with imaging probes likely reflect the dynamics of CB_1_ in neurons as both probes in live neurons displayed a spatial pattern similar to that of CB_1_ antibody in fixed neurons. Previous studies have shown that GPCRs form homodimers, heterodimers, or oligomers to affect their downstream signaling pathways ^28, 29, 30, 31^.

However, uncovering the cellular structure of GPCRs has proven to be a challenging task, as several prior studies did not observe the semi-periodic organization of CB_1_ as we did ^11, 27^. An early study using STORM to characterize the CB_1_ structure in brain tissue did not observe any sign of organized CB_1_ pattern in axons. This could be due to the use of different antibodies, but most likely it was because they did not focus on CB_1_ in the axonal shaft region, but instead focused on axonal boutons ^11^. Focusing on cultured neurons, Zhou et al. found that CB_1_ displays a non-periodic structure without the application of WIN, an agonist of CB_1_, and becomes periodic upon the administration of WIN ^27^. They concluded that the MPS acts as a dynamically regulated platform for GPCR-related signaling. Our results using the same antibody used in Zhou’s experiments showed both periodic and non-periodic hotspots in both culture neurons and native brain tissue. We observed the periodicity in live neurons to avoid artifacts caused by fixation procedures. Further, we found that active CB_1_ may behave differently than native ones, as CB_1_ displayed less dynamic and more confined movements upon WIN application. Actually, Zhou’s data implied that there was weak CB_1_ periodicity and colocalization with spectrin at ~190 nm before agonist stimulation ^27^. The periodic distribution without application of WIN could be explained the fact that the constitutive released ligand, such as 2-AG, may activate CB_1_ to certain extent and thus leads to the periodic pattern of CB_1_. Further studies will be required to answer the mechanism and significance of these constitutive periodic pattern of CB_1_.

In conclusion, we found that CB_1_ is distributed in axons as organized hotspots separated by approximately 190 nm, in a similar spatial distribution to the MPS. Especially, CB_1_ tends to be more organized as periodic hotspots upon agonist application suggesting that active CB_1_ associate more strongly with the MPS. When a GPCR is activated by an agonist, it increases the kinetics of interaction with G proteins and conducts downstream signal transduction within hotspots, which are usually confined to the cytoskeletal and clathrin-forming grids ^32^. Our results suggest that active CB_1_ is clustered into hotspots where G proteins or β-arrestin can easily collide with and form a relatively stable interaction with intracellular signaling components to increase signaling efficacy. Our results also indicate that CB_1_ are anchored to the MPS, likely by ankB, forming a fundamental structural unit that may be important for the proper function of neurons. These units likely form the structural basis for hotspots where signals transfer from extracellular to intracellular compartments. Our observation of periodic hotspots of CB_1_ along axon shafts, and the role of the cytoskeleton in CB_1_’s intracellular signaling, indicate a new horizon for the study of structural-functional interaction in neurons.

## Methods

### Animals

C57BL/6 mice, Sprague-Dawley rat, CB_1_ knockout mice (Biocytogen) and Kunming (KM) mice (for knockout mice generation) were used in this study. Animals were housed under a 12 h light/dark cycle at a room temperature of 22 ± 1°C with food and water available *ad libitum*. All experimental procedures were approved by the Institutional Animal Care and Use Committee of ShanghaiTech University, China.

### Generation of *Cnr1* knockout mouse

For *Cnr1* gene targeting, two sgRNAs were designed to target either an upstream or downstream region of its coding sequence by the CRISPR design tool (http://crispr.mit.edu) and screened for on-target activity using UCA^TM^ (Universal CRISPR Activity Assay, Biocytogen). PCR amplification was performed to add the T7 promoter sequence to the Cas9 mRNA and sgRNAs DNA template and then the T7-Cas/sgRNA PCR products were gel purified. They were used as the template for *in vitro* transcription with the MEGAshortscript T7 kit (Cat. No. AM1354, Life Technologies). The Cas9 mRNA and sgRNAs products were purified with MEGAclear kit (Cat. No. AM1908, Life Technologies) and eluted with RNase-free water.

C57BL/6 female mice and KM mouse strains were used as embryo donors and pseudo-pregnant foster mothers, respectively. Superovulated female C57BL/6 mice (3-4 weeks old) were mated to C57BL/6 stud males, and fertilized embryos were collected from the ampullae. Different concentrations of Cas9 mRNA and sgRNAs were mixed and co-injected into the cytoplasm of one-cell stage fertilized eggs. After injection, surviving zygotes were transferred into oviducts of KM albino pseudo-pregnant females and allowed to develop to term.

Mutant mice were genotyped to ensure the deletion of target CB1 segment. To mitigate off-targets effects, mutant mice were crossed into C57Bl/6 for 5 generations before being used for experimental purposes.

### Primary culture of rat hippocampal neurons

Sprague-Dawley rats of either sex at P0 were used for culturing rat hippocampal neurons. Hippocampi were isolated and digested with papain (1mg/mL; Sigma, P3125) at 37°C for 30 min. The digested tissues were washed with DMEM solution (Hyclone, SH30243.FS) three times and then transferred to culture medium containing Neurobasal medium (Thermo Fisher Scientific, 21103049) supplemented with 2% (vol/vol) B27 supplement (Thermo Fisher Scientific, 17504044) and 1% (vol/vol) Glutamax (Thermo Fisher Scientific, 35050–061). The tissues were gently triturated until no chunks of tissue were left. Dissociated cells were then counted and plated onto poly-D-lysine–coated 12-mm coverslips (12-545-80, Fisher Brand) or 29mm Glass (#1.5 cover glass) bottom dishes (D29-20-1.5-N, Cellvis). Neuronal cultures were maintained in the culture medium in a humidified atmosphere with 5% CO_2_ at 37°C. One-half of the medium was changed every 3 days to maintain neuron viability.

### Fluorescence labelling of neurons

Cultured neurons were fixed with 4% (w/v) paraformaldehyde in phosphate buffered saline (PBS) for 15 min at 14 day *in vitro* (DIV 14). After a complete wash with PBS, the samples were then permeabilized and blocked in a blocking buffer (10% v/v donkey serum, 0.2% v/v Triton X-100 in PBS) for 1 h at room temperature, and subsequently stained with one or two primary antibodies in the incubation buffer (1% donkey serum, 0.1% Triton X-100 in PBS) overnight at 4°C. The samples were washed three times and then stained with secondary antibodies as described above in the incubation solution for 1 h at room temperature. Neuron samples were mounted with ProLong Gold (Life Technology, for STED imaging) or Vectashield (Vectorlabs, for SIM imaging) antifade mounting media.

### STED Imaging

STED images were obtained using the Leica TCS SP8 STED 3X microscope, equipped with white light pulse laser (WLL2), STED laser (592 nm, 660 nm), oil immersion 100x / NA 1.4 objective lens (HC PL APO CS2, Leica), hybrid detectors with a high sensitivity, and TCS SP8 time-gated system. The STED depletion laser was co-aligned with the excitation laser, and selectively deactivated the excited fluorophores surrounding the focal point by stimulated emission, leaving the remaining fluorescence in the center to be detected. This approach allowed an increased resolution up to 50 nm by shrinking the point-spread function (PSF) of the microscope. Images were acquired in both confocal mode and STED mode with 1024 × 1024 pixel dimensions. Acquisition settings such as laser power, image size, pixel dwell times, line average, frame accumulation and time-gating interval (1-6 ns post-pulse time window) were optimized for achieving the best image quality. Deconvolution of STED images were performed by Huygens software (Scientific Volume Imaging) with the Huygens classical maximum likelihood estimation (CMLE) deconvolution algorithm and theoretical Point Spread Functions. Images were deconvolved with the deconvolution wizard. Before deconvolution, we checked parameter settings in the software and confirmed the restore of the acquisition parameters.

### Live SIM imaging the structure of CB_1_ in neurons

SIM images were obtained with the GE DeltaVision OMX microscope, equipped with a 568-nm laser (coherent), an oil immersion 60x / NA 1.42 objective lens (PlanApo N, Olympus), and a scientific CMOS camera (Acquisition pixel size: 82 nm at 60x objective; PCO edge). 3D-SIM mode was used with a fixed Z interval of 0.125 μm, and 15 images were taken with 3 angles and 5 phases for each Z-section. Images were taken in fast 286 MHz mode. To obtain images with minimized spherical aberration and optimized illumination contrast, immersion oil with different refractive indexes was systematically optimized for each sample. The raw images were reconstructed by the OMX SI reconstruction tool available in softWoRx (GE). Channel-specific OTF files, channel-specific K0 angles, Wiener filter constant (0.001), and bias offset (65) were used during the reconstruction.

CB_1_-RFP was transfected in the cultured neurons 24 h before being imaged under the 3D-SIM mode at different time points. The cells were incubated with 0.01M PBS at the 37℃ and supplied with 5% CO_2_ mixed with 95% air during the imaging process. The exposure times were set as 7-10 ms per frame and acquisition interval was 1 or 3 min. An ultimate focus system was used to maintain the sample Z position, regardless of the mechanical and thermal changes during the acquisition. After SIM reconstruction, the images were processed in the FIJI software. The image drift during acquisition was first corrected and then at each time point, we calculated the maximum projections of image and adjusted the brightness and contrast. Fiji TrackMate was then used to analyze the dynamics of CB_1_ hotspots in these images^33^. In TrackMate, the difference of Gaussian (DoG) detector was used with an estimated spot diameter of 80 nm and an appropriate fluorescence intensity as the threshold to detect all the individual CB_1_ hotspots. The Simple Linear Assignment Problem (LAP) tracker with a linking maximum distance of 200 nm, a gap-closing maximum distance of 200 nm, and a gap-closing maximum frame gap of 2 was used for tracking the cells through the time course images.

### In situ PLAs

Cells stably expressing CB_1_ and transfected with the corresponding cDNA were grown on glass coverslips and were fixed in 4% paraformaldehyde for 15 min, washed with PBS containing 20 mM glycine, permeabilized with the same buffer containing 0.1% Triton X-100, and successively washed with PBS. Interactions between CB_1_ and cytoskeleton proteins were detected using the Duolink *in situ* PLA detection Kit (Sigma) following the instructions of the supplier. To detect CB_1_-cytoskeleton protein interaction, a mixture of equal amounts of anti-CB_1_ antibody directly linked to a plus PLA probe and cytoskeleton protein antibody (like ankyrin B or βII-spectrin) directly linked to a minus PLA probe was used. The PLA probe was linked to the antibodies following the instructions of the supplier. Cells were mounted using the mounting medium with DAPI. The samples were observed in a Nikon confocal microscope equipped with an apochromatic 60X oil-immersion objective (N.A. 1.4), and a 405 nm and a 561 nm laser line. For each field of view, a stack of two channels (one per staining) and 10 to 15 Z stacks with a step size of 1 μm were acquired. Images were opened and processed with Fiji software. The ratio r (number of red spots/number of cells containing spots) was determined considering a total of 300 - 600 cells from 6 - 10 different fields.

### Western blot

Primary neurons were either treated or not treated with the indicated ligands for the times noted, rinsed with ice-cold PBS, and lysed by the addition of 100 μl of ice-cold RIPA lysis buffer (Beyotime) with protease inhibitor cocktail (Roche). Cellular debris was removed by centrifugation at 13,000 g for 15 min at 4°C, and the amount of protein was quantified by BCA protein assay kit (Pierce). Protein samples were separated on 10% SDS-PAGE gels (Bio-Rad) and transferred onto PVDF (polyvinylidene fluoride) membranes (Millipore, Billerica, MA, USA). The membranes were blocked with 5% non-fat dry milk for 2 h at RT and incubated overnight at 4°C with rabbit anti-phospho-ERK1/2 (1:2000, Cell Signaling), anti-phospho-Ser473-Akt (1:2000, Cell Signaling), anti-ERK1/2 (1:2000, Cell Signaling), and anti-AKT (1:2000, Cell Signaling) antibodies. The membranes were then incubated with HRP-conjugated secondary antibody (1:1000; Pierce) for 2 h at room temperature. Signals were visualized using enhanced chemiluminescence (ECL, Pierce, Rockford, IL), and captured by the ChemiDoc XRS system (Bio-Rad Laboratories, CA). Phosphorylated ERK1/2 or phosphorylated Akt levels were normalized for differences in loading using protein band intensities for total ERK1/2 or AKT.

### Immunoprecipitation

Following tetracycline induction, CB_1_-CHO cells were washed with ice-cold phosphate-buffered saline (PBS) and suspended in immunoprecipitation (IP) buffer containing (in mM): 50 Tris-HCl, 120 NaCl, 0.5% Nonidet P-40 and protease cocktail (PH=7.5). The lysate was sonicated, centrifuged at 13,000 rpm for 20 min at 4°C, and the resulting supernatant was incubated with the rabbit anti-CB_1_ (CST, 93815) antibody for 20 min at 4°C. Immuno-complex was incubated with Protein A-Magnetic beads overnight on a rotating wheel at 4°C from the precleared supernatant with anti-CB_1_ antibody covalently coupled to Protein G-Magnetic beads. The pellet was then washed 5◻times◻in wash buffer containing (in mM): 20 Tris-HCl, 100 NaCl, 1 EDTA, 0.5% Nonidet P-40 (PH=8.0). The beads were to wash away any proteins non-specifically bound to the beads. The immunoprecipitates were mixed with the loading buffer and resolved by SDS-PAGE. Western blots were performed with relevant antibodies. Rabbit IgG was used as a negative control.

### Mass spectra in brain tissue

#### Mouse brain tissue preparation

Preparation of mouse brain membrane fractions was performed according to a previous study^34^. Briefly, 6 brain regions (olfactory bulb, cerebral cortex, cerebellum, hippocampus, midbrain, and spinal cord) were obtained from 9-week-old C57BL/6 wild-type male mice. Brain regions from three mice were pooled and homogenized in the buffer of 300 mM sucrose, 0.5% BSA, 100 mM EDTA, 30 mM Tris/HCl, pH 7.4 with protease inhibitor (Roche). Crude membrane fractions were isolated from the homogenate by ultra-centrifugation at 160,000 g at 4°C for 1 h. The membrane pellet was solubilized in 4% SDS and 100 mM DTT in 100 mM Tris/HCl, pH 7.6, denatured and reduced at 95°C for 3 mins. Protein concentration was determined using BCA assay. For each brain region, protein sample preparation was conducted in duplicate.

#### Protein digestion

The SDS-assisted digestion of membrane proteins was performed according to methods described previously^35^. Briefly, 50 μg of protein was diluted in 8 M urea, 50 mM NH_4_HCO_3,_ and exchanged to the same buffer using the 30 KDa MWCO centrifugal filter unit (Satorious, Germany) by centrifugation at 13,000 g for 20 min. The following centrifugation steps were performed under the same conditions. Subsequently, 100 μl of 50 mM iodoacetamide in 8 M urea, 50 mM NH_4_HCO_3_ was added and incubated at room temperature in darkness for 30 min, then followed by centrifugation. The concentrate was diluted with 200 μl 50 mM NH_4_HCO_3_ and centrifuged again, this step was repeated twice. Proteins were digested with trypsin (Promega, Madison, USA) at an enzyme-to-protein ratio of 1:100 (w/w) at 37°C for 3 h, followed by the addition of trypsin at 1:50 (w/w) and incubation at 37°C overnight. After acidification, the protein digest was desalted with C18-SepPak columns (Waters, Milford, USA) and lyophilized under vacuum.

#### NanoLC-MS/MS analysis

The nanoLC-MS/MS analysis was conducted on an EASY-nLC 1000 connected to Orbitrap Fusion mass spectrometer (Thermo Fisher Scientific, USA) with a nano-electrospray ionization source. The eluted peptides were separated on an analytical column (200 mm × 75 μm) in-house packed with C18-AQ 3 μm C18 resin (Dr. Maisch, GmbH, Germany) over a 130-min gradient at flow rate of 300 nl/min. For a pooled sample from all six brain region protein digests, a data-dependent (DDA) acquisition method was first employed with the following parameters: resolution of 60,000 used for survey scans; the mass range pf 300 - 1700 m/z; an AGC target value of 4E5; and maximum ion injection time of 50 ms. Up to 12 dynamically chosen and most abundant precursor ions were fragmented. The MS/MS scans were acquired at an Orbitrap resolution of 30,000 (AGC target value 1E5, maximum ion injection time 50 ms).

In order to achieve the accurate quantification of selected proteins in the membrane fractions, we developed parallel-reaction-monitoring (PRM) MS assays for all proteins of our interest based on the protein identification results from the DDA experiment^36^. The PRM acquisition method started with a full scan event followed by targeted MS/MS for specific peptides from the proteins of interest. Major parameters for the MS/MS event in Orbitrap were: resolution of 30,000; an AGC target value of 2E5; and maximum injection time of 100 ms. Peptide precursor ions in different brain region protein digests were monitored in the PRM assay by scheduling an inclusion list of each precursor with an isolation window of 1.6 m/z and retention time shift of 2 min.

#### MS data processing

Mass spectra from the DDA experiment were processed using Proteome Discoverer 2.1 against the Uniprot mouse sequence database. The search parameter included cysteine carbamidomethylation as a fixed modification and oxidation of methionine as variable modification. Precursor ion mass tolerance was set to 10 ppm and fragment ion tolerance was 0.02 Da. Trypsin was set as the specific enzyme and two missed cleavages were allowed. The required false discovery rate (FDR) was set to 1% at the peptide and protein level.

For PRM data analysis, the Skyline software (v3.7.0) was used for targeted peptide quantification with settings specified by the software instruction^37^. Only b- or y-product ions with m/z values greater than the precursor were selected to quantify the peptides. All the transitions were validated using the mProphet algorithm in Skyline advanced peak picking model that restricts the false discovery rate to <1%. The peptide quantification was derived from the sum of the peak areas of three to six product ions for selected peptides. Protein intensity was based on the summed MS responses of one to three unique peptides of the corresponding protein.

### Quantification and Statistical Analysis

All image data were first processed with Fiji software (National Institutes of Health). Images were resized with the Bicubic interpolation and the brightness and contrast were linearly adjusted. To quantitatively analysing the distribution properties of CB_1_, segmented lines across the structures were drawn, the intensity profiles along the lines were measured and further analysed in Matlab (MathWorks, Inc.). Individual fluorescence peaks were found and the distance between the neighbouring peaks were calculated and pooled together. For distribution pattern analysis, autocorrelation and cross correlation were performed on the fluorescence intensity profiles. The correlation curve was pooled and averaged from many randomly selected lines. All intensity, distance and correlation data were plotted using Graphpad prism (Graphpad Software, Inc.), and all figure layouts were composed in Illustrator (Adobe Systems, Inc.).

Results were reported as mean ± SEM. Statistical analysis of the data was performed using a student t test, one-way ANOVA. Statistical significance was set at p < 0.05.

## Author Contributions

G.Z. initiated, managed and supervised the project. G.Z., and H.L. conceived and designed the experiments. H.L., Y.J., T.C., L.S., D.M., T.H., performed most of the experiments and data analysis. Z.L., D.G.D., G. T., Y.Z., W.S. and Z.J.L. the authors contributed to data analysis and interpretation. G.Z. wrote the manuscript with contributions from all of the authors.

## Acknowledgements

We thank the Bioimaging Core Facilities of the iHuman Institute and the animal facility of National Center for Protein Science for their support. We thank Dr. Yulong Li from Peking University for providing the GPCR probes. This work was supported by the National Key Research and Development Program of China (2016YFC0905900 (G.Z.), 2017YFC 1001300 (G.Z.)), the National Natural Science Foundation of China (81970878 (G.Z.), 31771130 (G.Z.), 81861128023(G.Z.)), the 2015 Thousand Youth Talents Plan of China (G.Z.).

---

**Supplementary Figure 1.**
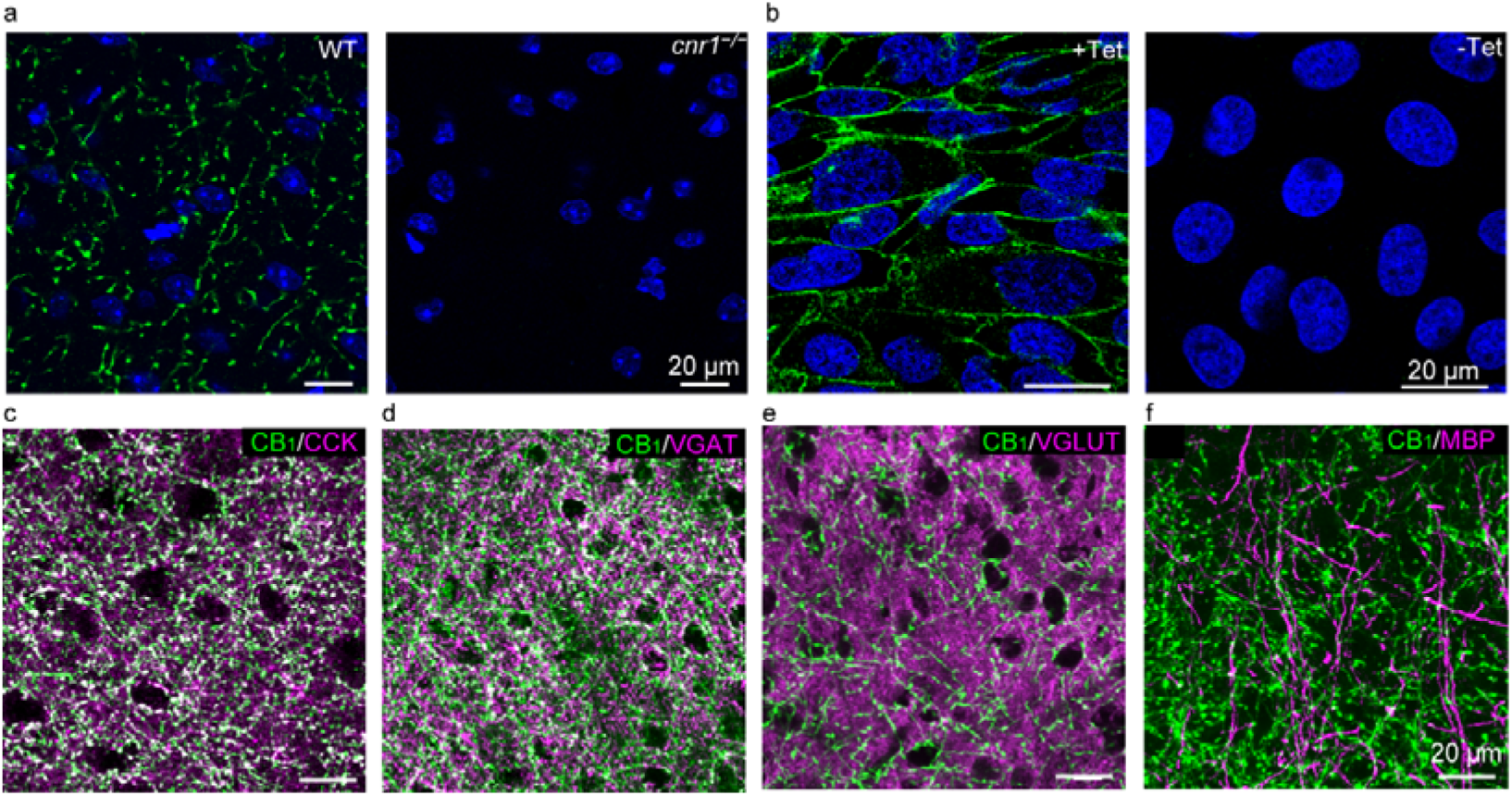
Confocal imaging reveals the antibody specificity of CB_1_ **a** Immunostaining of CB_1_ (green channel) in wild-type mouse (left panel) and *cnr1*^−/−^ mouse (right panel). **b** Immunostaining of CB_1_ with the antibody used in a CB_1_-CHO cell line with (left) and without (right) tetracycline application. **c-f** Representative fluorescent images of: (**c)** CCK; (**d)** VGAT; (**e)** VGLUT; and (**f)** MBP, all colocalized with CB_1_.

**Supplementary Figure 2.**
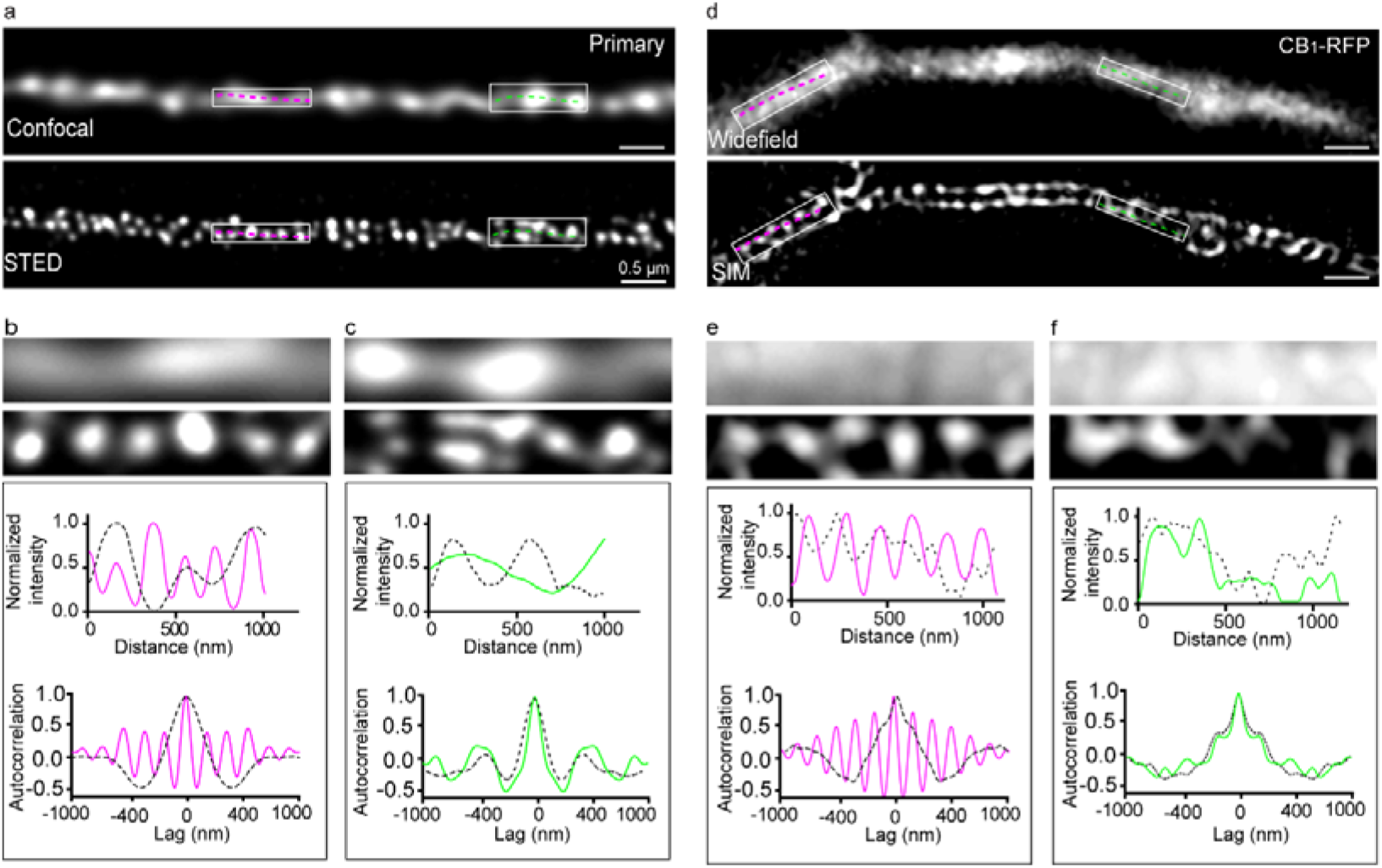
The semi-periodic hotspots of CB_1_ in cultured neurons **a** Representative confocal and corresponding STED image of CB_1_ labeled by antibody in culture hippocampal neurons. **b-c** Corresponding magnification of the two boxed regions in **a** (top) with intensity and autocorrelation analysis along the lines in the box regions below. **d** Representative wild field and corresponding SIM images of CB_1_ labelled CB_1_-RFP in culture hippocampal neurons. **e-f** Corresponding magnification of the two boxed regions in **d** (top) with intensity and autocorrelation analysis along the lines in the box regions below.

**Supplementary Figure 3.**
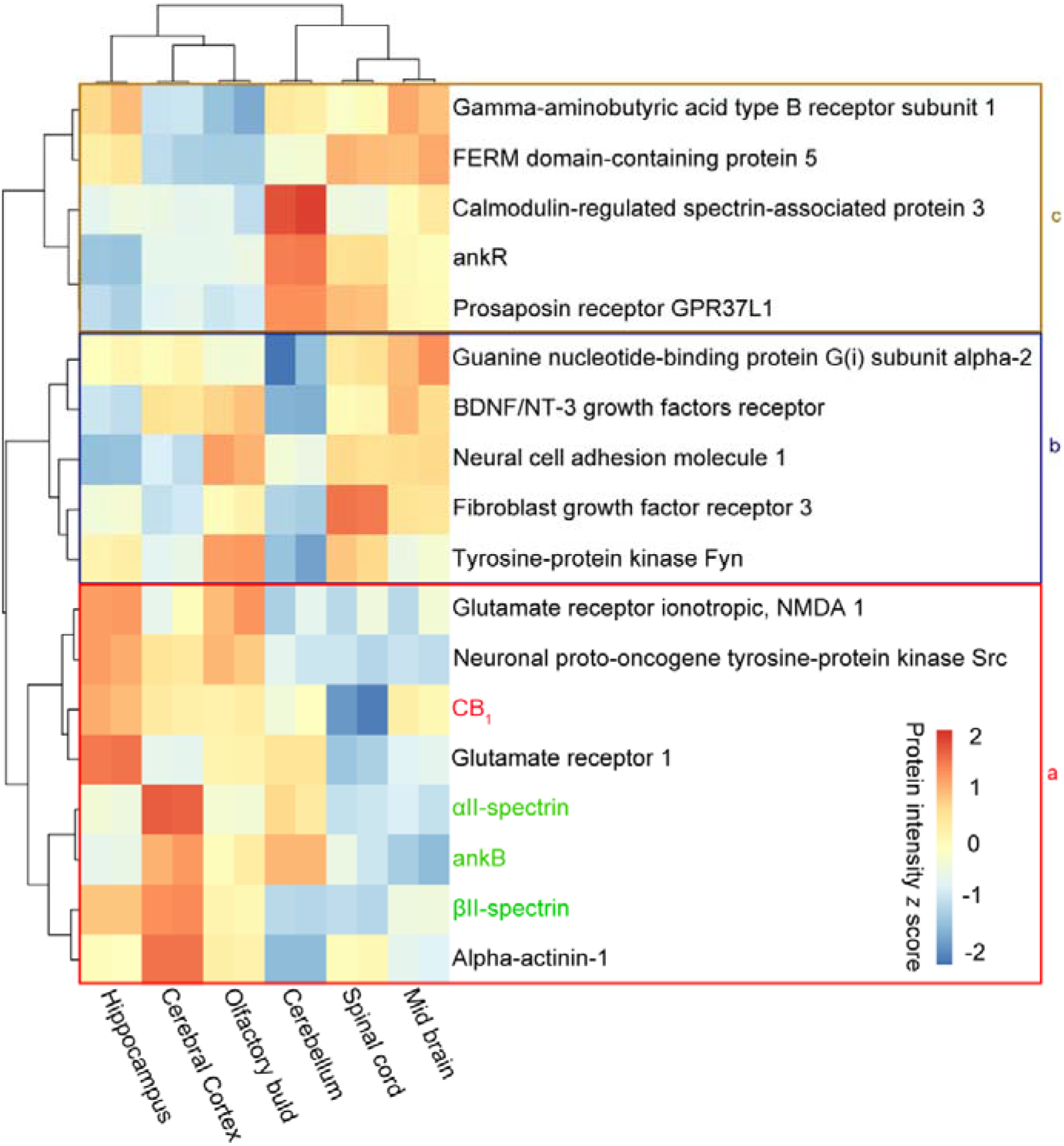
CB_1_ and associated proteins revealed by MS Heat map of three groups of selected protein expressed across different brain regions (n = 2 for each region). Heat map of z-scored protein abundances of the differentially expressed proteins (determined with PRM MS assays) after unsupervised hierarchical clustering revealed the correlation on the expression level of CB_1_ to cytoskeleton-related proteins **(a)**, receptors, signaling molecules **(b)**, and other less likely related proteins **(c).**

**Supplementary Figure 4.**
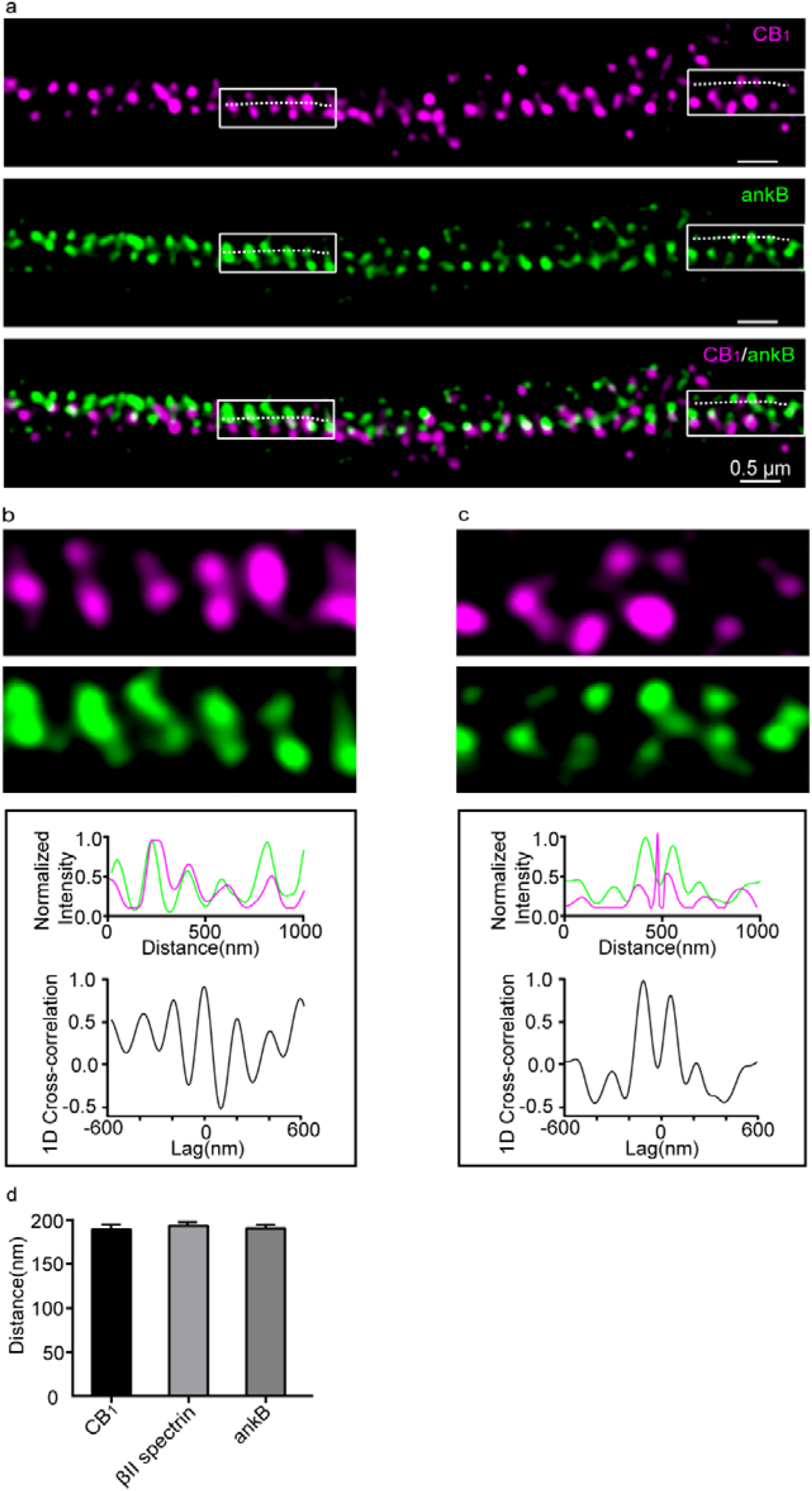
Spatial relationship between CB_1_ and ankB revealed by STED **a** Two-color STED image of CB_1_ (magenta) and ankB (green) in the axons of culture neurons. The white boxes represent regions enlarged in **b** (left box) and **c** (right box). **b-c** Enlarged regions from **a** (top). Normalized intensity and 1D cross-correlation analyses between the distributions of CB_1_ and ankB from white lines in **a** (bottom). **d** Comparison of the periodic spacing between CB_1_ and cytoskeleton components in the primary neurons. Data are mean ± s.e.m. (n = 3 biological replicates; 70-120 axonal regions were examined per condition). P = 0.9226, not significant (p > 0.5), one-way ANOVA. Actual spacing (from left to right): 188 ± 5 nm, 192 ± 1 nm, 188 ± 1 nm.

**Supplementary Figure 5.**
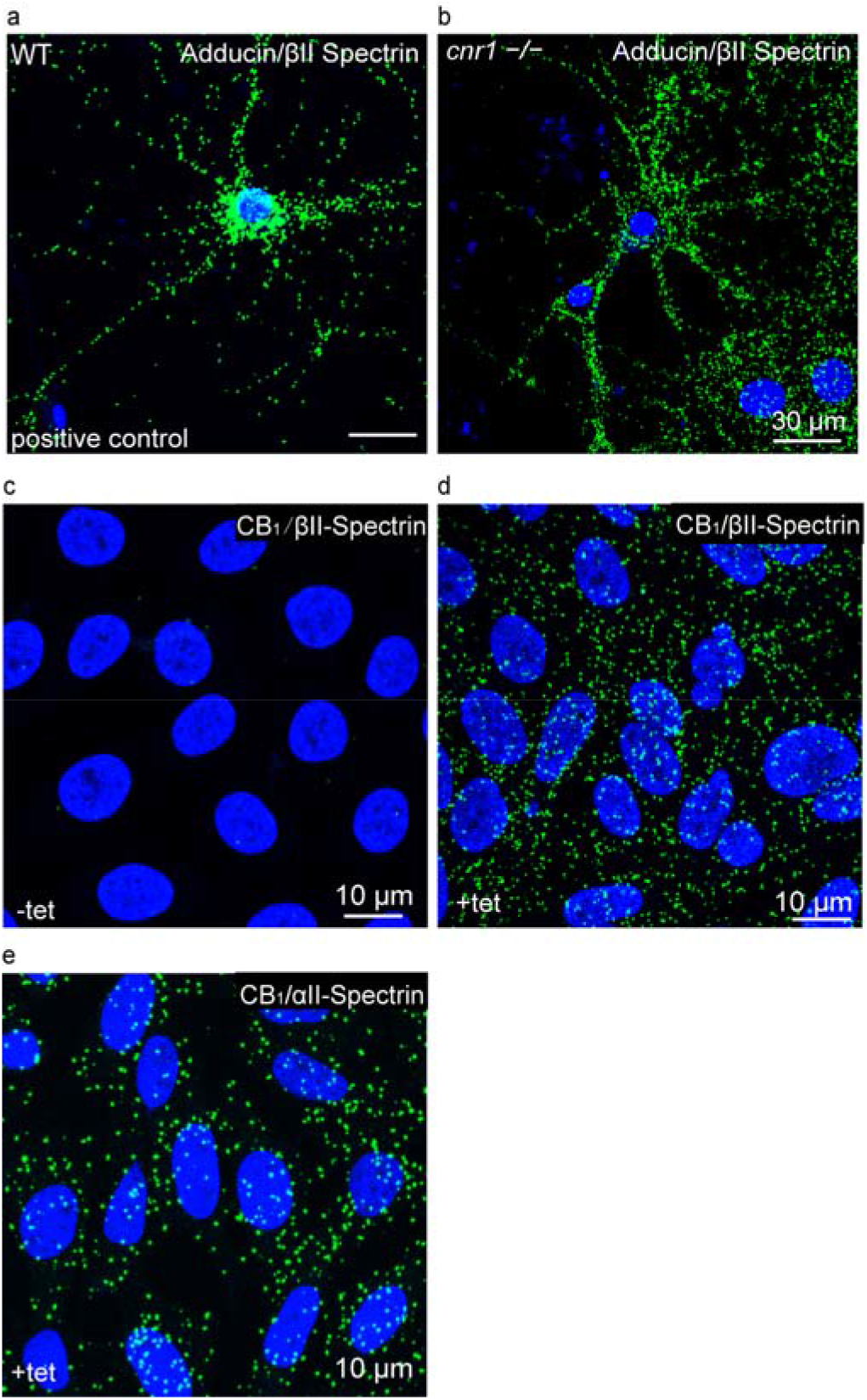
PLA can detect the proteins within a short distance. **a-b** PLA signals were detected in the cultured neurons of both WT (**a**) and *cnr1*^−/−^ mouse (**b**) with antibody of CB_1_ and adducin, which have certain interactions. **c-d** PLA analyses of CB_1_ and βII-spectrin in CB_1_-CHO cells in the absence (**c**) and presence (**d**) of tetracycline application. **e** PLA results of CB_1_ and αII-spectrin in CB_1_-CHO cells with tetracycline application.

**Supplementary Figure 6.**
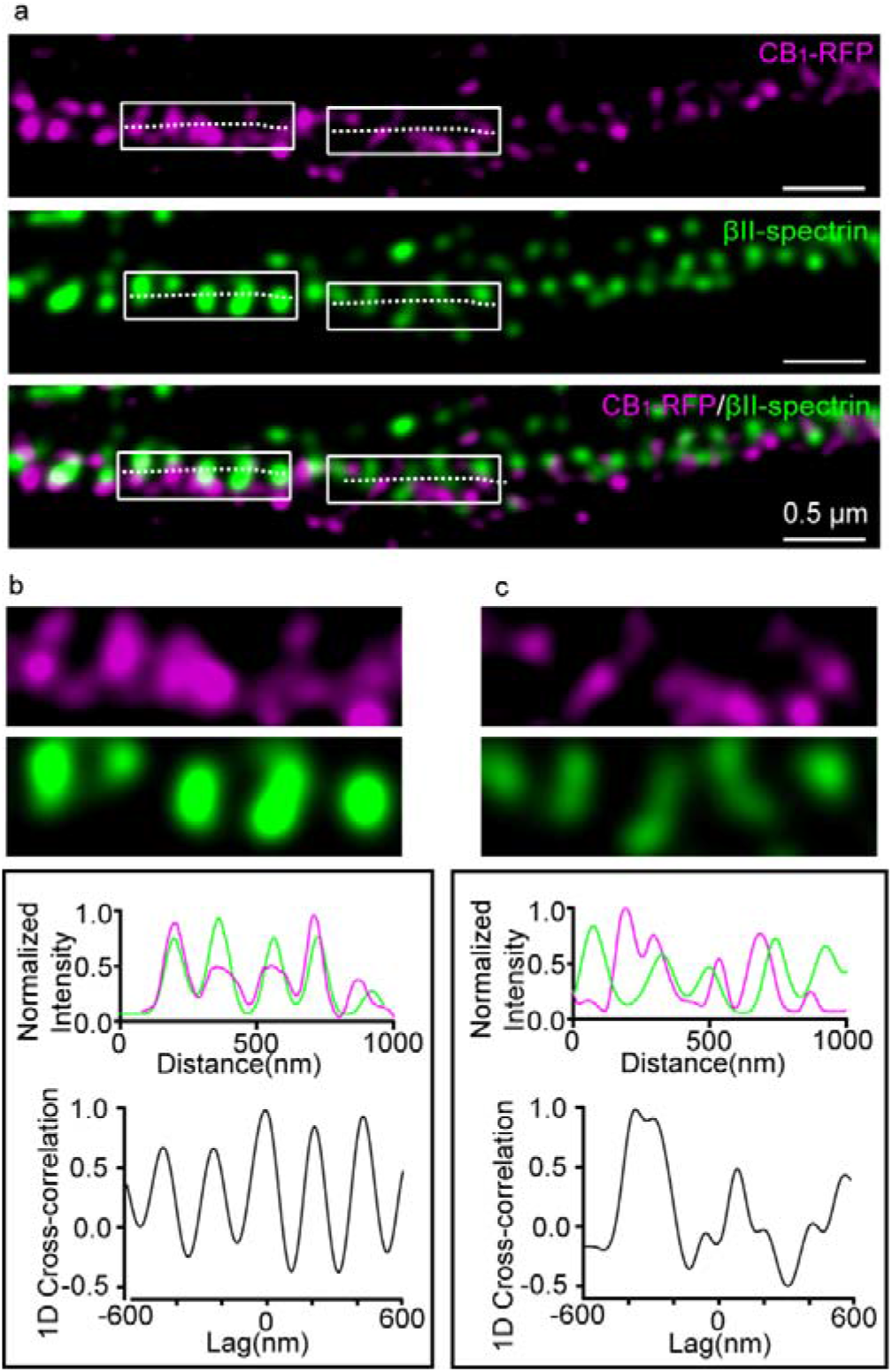
Spatial relationship between CB_1_-RFP and βII-spectrin revealed by STED **a** Two-color STED image of CB_1_-CFP (magenta) and βII-spectrin (green) in the axons of cultured neurons. The white boxes represent regions enlarged in **b** (left box) and **c** (right box). **b-c** Enlarged regions from **a** (top). Normalized intensity and 1D cross-correlation analyses between the distributions of CB_1_-CFP and βII-spectrin from white lines in **a** (bottom).

**Supplementary Figure 7.**
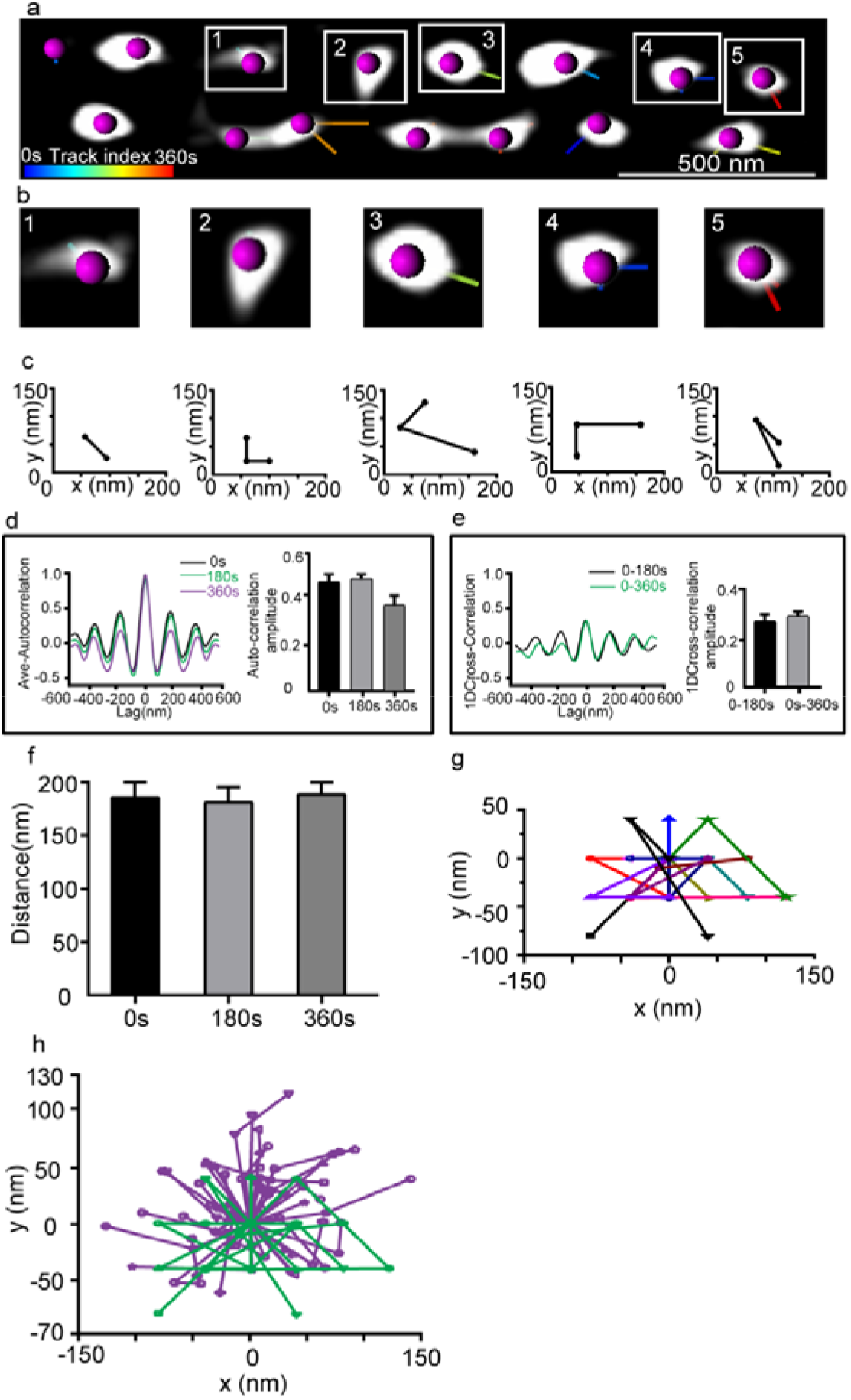
The dynamics of CB_1_ hotspots revealed by live SIM imaging **a** Representative live image of transfected CB_1_-RFP in the primary neuron (DIV 9-12) of the SD rat acquired by SIM taken at an interval of 3 mins. Each CB_1_ hotspot was marked with a purple dot, and its location at each individual time point is connected by a line. The color of line indicates the trace index. **b-c** The five CB_1_ hotspots shown in **a** and their relative locations. Displacements are around 60-70 nm between neighboring time points. **d** Averaged autocorrelation analysis of CB_1_ distributions at different time points with the autocorrelation amplitude shown on the right. There was no statistical difference between time points. P = 0.88, not significant (p>0.5), one-way ANOVA. Actual autocorrelation amplitude (from left to right): 0.42 ± 0.07, 0.42 ± 0.05, 0.37 ± 0.08. **e** Averaged cross-correlation analysis between neighboring frames (0s-180s, 0s-360s) showed similar distribution properties (left) and amplitude (right). P = 0.84, not significant (p>0.5), unpaired Student’s *t* test. Actual cross-correlation amplitude (from left to right): 0.25 ± 0.09, 0.27 ± 0.04. **f** CB_1_ spacing across time points. P = 0.9163, not significant, one-way ANOVA. Actual spacing (from left to right): 192 ± 1 nm, 190 ± 1 nm, 193 ± 1 nm. **g** Dynamics of the individual CB_1_ hotspots over time. **h** Overlay of the dynamics of the individual CB_1_ hotspots over time taken at interval of 1 min (purple) or 3 min (green). The moving distance did not increase during imaging, suggesting that individual hotspots displayed a confined dynamic around its original point. Data in **d**, **e** and **f** are mean ± s.e.m (n = 3 biological replicates; 70-120 axonal regions were examined per condition).

**Supplementary Figure 8.**
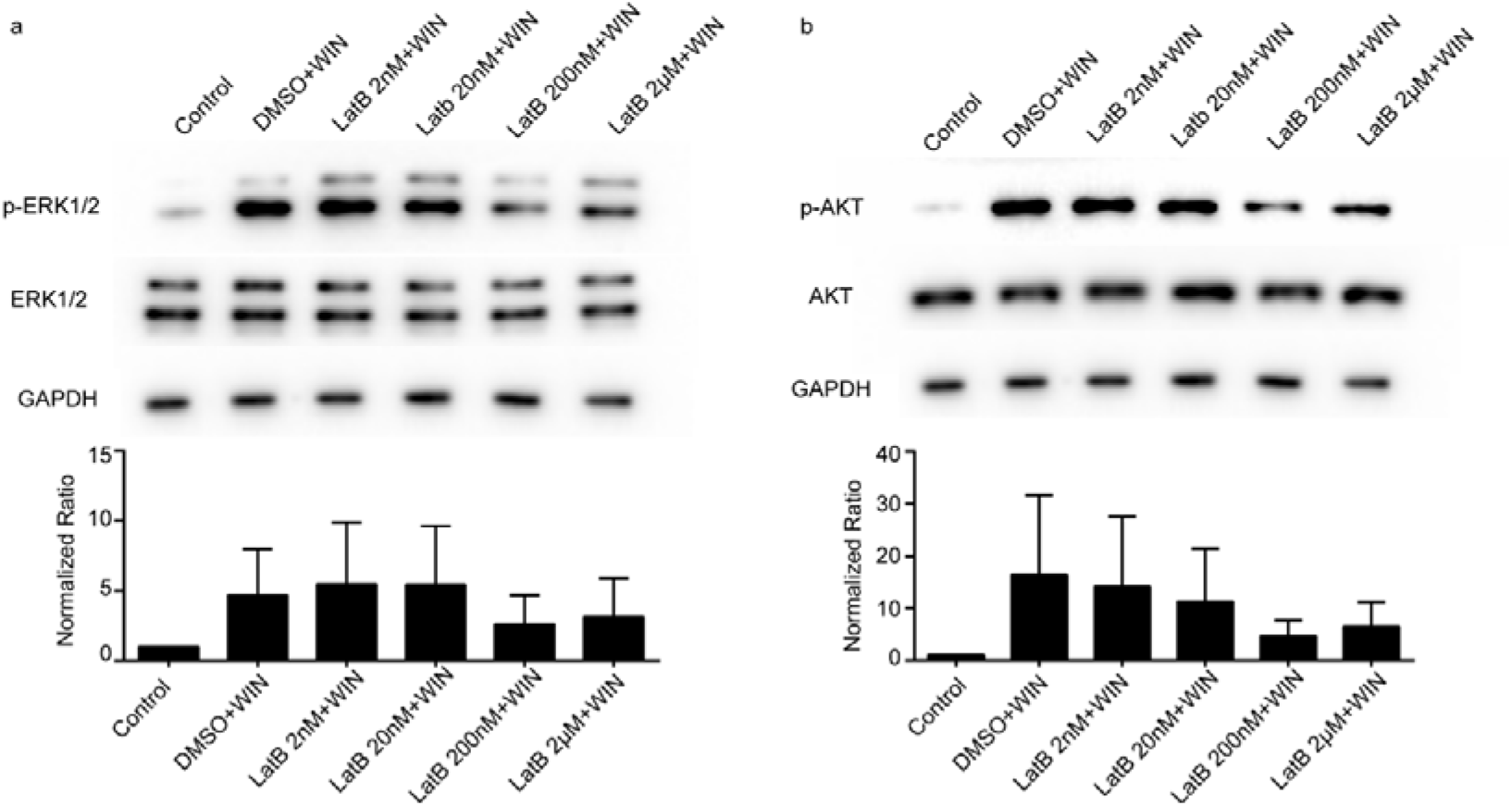
CB_1_ signaling is related to the cytoskeleton **a** Western blot analysis for ERK1/2 and phosphorylated ERK1/2 in whole-cell lysates from primary neuron cells treated with different concentrations of latB. Western blots are representative examples from at least 3 independent biological replicates. **b** Western blot analysis for Akt and phosphorylated Akt in whole-cell lysates from primary neuron cells treated with different concentrations of latB.

